# FishCODE: a web-based information platform for comprehensive omics data exploration in fish research

**DOI:** 10.1101/2024.09.25.614839

**Authors:** Heng Li, Wanting Zhang, Keyi Ren, Hong Yang, Lei Zhang, Waqar Younas, Yingyin Cheng, Yaping Wang, Mijuan Shi, Xiao-Qin Xia

## Abstract

In terms of the utilization of omics data, the current fish database analysis functions are primarily relatively simple tools at the transcriptional level, aimed at obtaining the co-expression levels of specified genes or the data visualization of multiple genes, and do not enable users to perform comprehensive omics data analysis. Furthermore, the gene-level information currently provided by these multispecies fish genomics databases is incomplete, and there is a lack of a comprehensive portal that can offer multidimensional genetic information. To address these challenges, we collected extensive multi-omics information on 35 fishes and established the primary comprehensive multi-omics data information platform for fish, FishCODE (http://bioinfo.ihb.ac.cn/fishcode). We have collected experimental background of dataset which pertaining to the target fishes, selected a range of datasets that encompass a broad spectrum of research areas, and downloaded the corresponding raw omics data from public repositories such as the Sequence Read Archive (SRA). Through a unified pipeline analysis, FishCODE contains 11,216 samples from 540 sets of genomic, transcriptomic, and methylomic datasets. These data encompass transcript structure and expression, gene methylation levels, protein domains, protein subcellular localization, protein interactions, best matched protein (Swiss-Prot), associated SNP site information (47,111,018), orthologous genes, phylogenetic tree and GO/KEGG annotations. To facilitate comparison, we annotated the experimental background data sets of the FishCODE, FishGET, PhyloFish, FishSED and FishSCT databases using the Fish Experimental Condition Ontology. Currently, the FishCODE database omics dataset includes 146 unique experimental condition words, 654 cumulative experimental condition words, and 13 species with rich experimental background (more than 20 unique FECO words). These data are 3.5 times (42), 8.3 times (74), and 6.5 times (2) those of the second-ranked databases respectively. We generated word cloud maps for the experimental condition vocabularies of FishCODE and FishGET, illustrating the superior richness of FishCODE’s experimental background.

## RESULT

Fishes constitute the largest group of vertebrates, accounting for more than half of all vertebrate species on Earth. Important aquaculture species like Atlantic salmon, cod, and large yellow croaker, contribute to over 16% of the global seafood supply for human consumption (Food and Agriculture Organization of the United Nations, 2020). Zebrafish (*Danio rerio*), medaka (*Oryzias latipes*), three spine stickleback (*Gasterosteus aculeatus*) and other fish species are vital model organisms in life science research. These organisms play a crucial role in disease modelling, drug screening, environmental toxicity assessment, and the study of various life processes, such as aging, damage repair, evolution, and genetic mechanisms. Due to the rapid development of high-throughput sequencing technologies and the extensive research value of fish itself, there has been a remarkable surge in multi-omics data in fish across various life science fields. The integrated analysis of these data provides a powerful means to analyze gene regulatory mechanisms and presents unprecedented opportunities for comprehensively understanding biological systems.

Currently, there exist specialized fish omics databases. However, these databases have not fully integrated and utilized the multi-omics data, which span a broad spectrum of research areas and encompass diverse experimental contexts. ZFIN (Sprague et al., 2001) is dedicated to the curation of zebrafish-specific data, with its RNA-Seq data primarily relying on external resources, such as Expression Atlas (Papatheodorou et al., 2020), offering a modest collection of 14 zebrafish RNA-Seq datasets that encompass a limited range of research areas. FishGET (Guo et al., 2023a) focuses on constructing accurate fish reference transcripts, encompassing data from 8 fish transcriptomes, predominantly those related to long non-coding RNAs (lncRNAs). This resource highlights economically important fish species such as grass carp and rainbow trout, while also providing a singular baseline dataset for the model organism zebrafish. FishSED (Guo et al., 2024) and FishSCT (Guo et al., 2023b) provide mainly zebrafish spatial transcriptome and single-cell transcriptome data. However, the datasets included in these resources are limited in the scope of research areas, due to the current scarcity of such data for fish species. PhyloFish (Pasquier et al., 2016) offer a baseline transcriptome dataset for 24 fish species. In the field of fish epigenomics, there exists a notable gap, as no high-throughput database is currently dedicated to fish methylation data, highlighting an area ripe for further research and development.

In terms of the utilization of omics data, the current fish database analysis functions are primarily relatively simple tools at the transcriptional level, aimed at obtaining the co-expression levels of specified genes or the data visualization of multiple genes (Fig. S1A), and do not enable users to perform comprehensive omics data analysis (Fig. S1B). Furthermore, the gene-level information currently provided by these multispecies fish genomics databases is incomplete, and there is a lack of a comprehensive portal that can offer multidimensional genetic information.

To address these challenges, we collected extensive multi-omics information on 35 fishes and established the primary comprehensive multi-omics data information platform for fish – FishCODE (http://bioinfo.ihb.ac.cn/fishcode). We have collected experimental background of dataset which pertaining to the target fishes, selected a range of datasets that encompass a broad spectrum of research areas, and downloaded the corresponding raw omics data from public repositories such as the Sequence Read Archive (SRA) (Table S1). Through a unified pipeline analysis, FishCODE contains 11,216 samples from 540 sets of genomic, transcriptomic, and methylomic datasets (Table S2) (data processing and other detail information are in the Supplementary Document). These data encompass transcript structure and expression, gene methylation levels, protein domains, protein subcellular localization, protein interactions, best matched protein (Swiss-Prot), associated SNP site information (47,111,018), orthologous genes, phylogenetic tree and GO/KEGG annotations. To facilitate comparison, we annotated the experimental background data sets of the FishCODE, FishGET, PhyloFish, FishSED and FishSCT databases using the Fish Experimental Condition Ontology (FECO, as detailed in Supplementary Document) (Table S3). Currently, the FishCODE database omics dataset includes 146 unique experimental condition words, 654 cumulative experimental condition words, and 13 species with rich experimental background (more than 20 unique FECO words). These data are 3.5 times (42), 8.3 times (74), and 6.5 times (2) those of the second-ranked databases respectively (Fig. S2A, S2B) (Table S3, S4). We generated word cloud maps for the experimental condition vocabularies of FishCODE and FishGET, illustrating the superior richness of FishCODE’s experimental background (Fig. S2C, S2D).

FishCODE’s omics analysis toolkit is extensive, featuring genome analysis tools such as single nucleotide polymorphism annotation, phylogenetic tree construction, and comparative genome browsing, as well as transcriptome (regular transcriptome, cross-species transcriptome, time-series transcriptome analysis) and epigenome (genome-wide methylation region analysis) analysis capabilities. These tools are intricately linked to the database’s wealth of omics data with diverse experimental contexts. Users benefit from a streamlined process that negates the need for preliminary research, data downloading, cleansing, or uploading. With a single mouse click, they can perform exploratory a comprehensive reanalysis of omics data (or in combination with user-uploaded data) (Table S4), yielding insights into candidate genes related to experimental condition variations, enriched biological pathways, and metabolic pathway dynamics, among other valuable information. In addition, FishCODE systematically collects and supplementally annotates (Gasteiger et al., 2001; Binder et al., 2014; Szklarczyk et al., 2023) molecular data on fish, and covers more extensive gene-related information than other multi-species fish omics databases (Table S4).

FishCODE provides a fuzzy search function for genetic information, which can provide a more comprehensive list of candidate genes (refer to “Fuzzy search function” in Supplementary Document) to help users to find the exact gene of interest. The Gene Details page offers comprehensive information about the gene, including its DNA, RNA, and protein dimensions, as well as its GO/KEGG annotations (Fig. S3). Taking the zebrafish *tp53* (gene id: ENSDARG00000035559) gene details page as a case study, FishCODE shows foundational information about the *tp53* gene at the DNA level, including its description, location, sequence, orthologous genes and more (Fig. S4). Additionally, it offers a visual representation of the *tp53* gene structure, associated single nucleotide polymorphisms (SNPs) (Fig. 1A), a heatmap showcasing the average methylation rates of the *tp53* gene within specified datasets (Fig. S5A), and a phylogenetic tree of orthologous genes (Fig. S6). Users can click on a row in the SNP information table to obtain detailed information including SNP genomic location, functional annotation, flanking sequences, and more (Fig. S7).

**Fig. 1.**
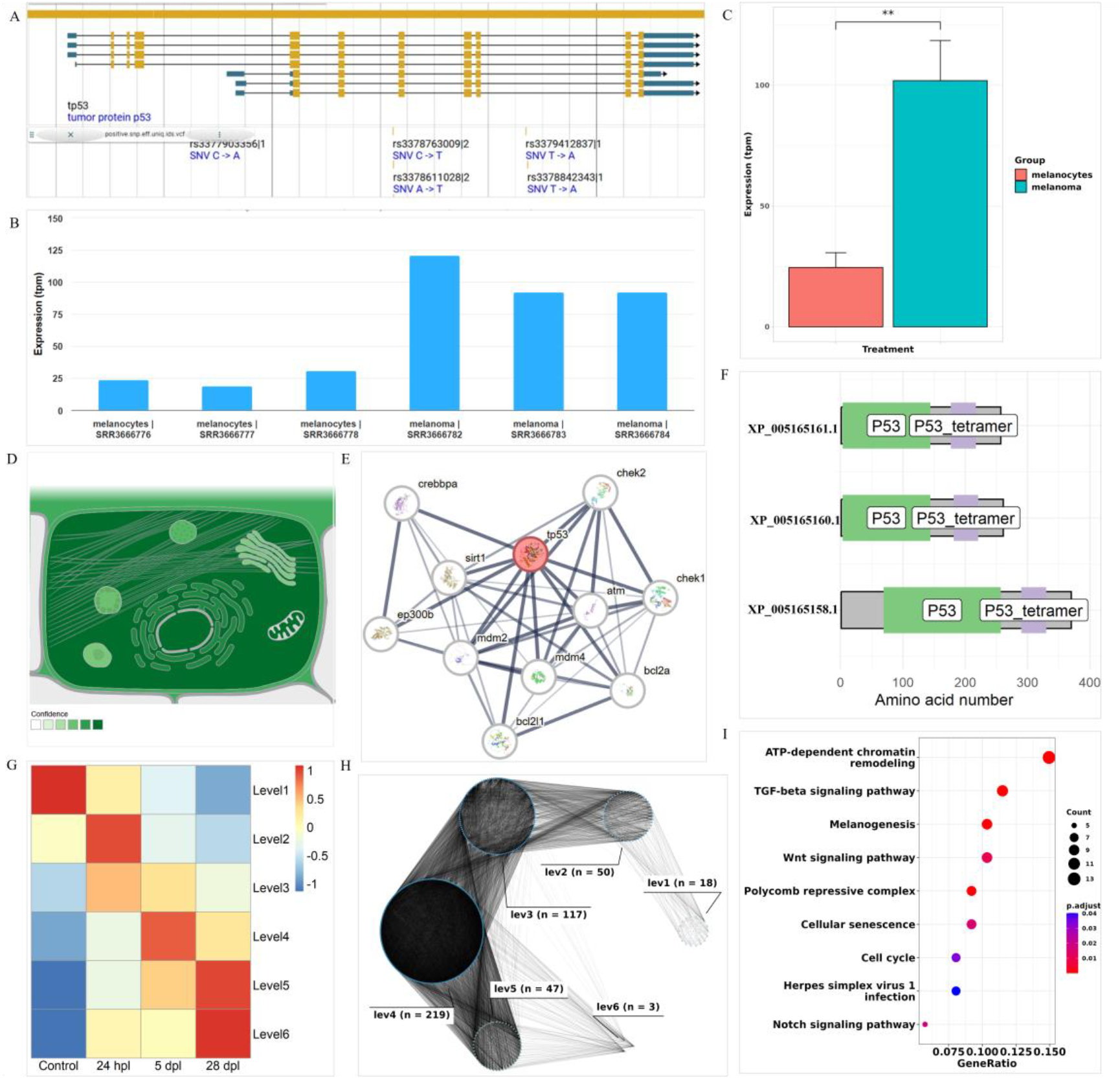
FishCODE’s features and content. A, Visualization of gene structure and variant sites. JBrowse2 (Diesh et al., 2023) plug-in enables users to visualize tp53 gene structure and SNPs sites, providing detailed information on gene structure and upstream and downstream regions. B, The bar plot of the expression level of the gene *tp53* (dataset: PRJNA325656). C, the bar plot of expression and statistics for gene *tp53* grouped by experimental treatment (dataset: PRJNA325656). This visualization is supported by Visual Omics (Li et al., 2023). D-F, Protein subcellular localization (darker organelle colors in protein subcellular localization indicate more reliable localization information), protein interactions, protein conserved structural domains of gene *tp53*. In addition, FishCODE also provides the above information on the most similar protein of the gene in the model species (human, mouse, rat, fruit fly, zebrafish), which users can freely switch to view (Fig. S13). G, Heatmap of time points and level correspondences. H, Network map of TF genes at different levels. I, KEGG enrichment results for level4-containing TF genes.

At the RNA level, users can obtain a bar plot of *tp53* gene expression information from any dataset (e.g., PRJNA325656, a dataset examining melanocytes and melanoma) (Fig. 1B) of interest, as well as detailed information about the corresponding samples. In addition, FishCODE enables users to retrieve gene expression levels and provide statistical differences between groups based on treatment, tissue, age, and gender groups. Continuing with dataset PRJNA325656 as an example, users can obtain statistical graphs grouped by experimental conditions (Fig. S8). The graph (Fig. 1C) reveals a significant upregulation of the *tp53* gene in melanoma cells, suggesting a potential correlation between *tp53* gene and melanoma which is supported by previous research (Hayward et al., 2017). At the protein level, users can access information and visualizations of subcellular locations, interactions, conserved structural domains (Fig. 1D, 1E, 1F), and the best matching proteins for the corresponding proteins of the *tp53* gene.

FishCODE encompasses an extensive toolkit for data analysis, featuring genome SNP annotation capabilities that can evaluate the potential impact of mutations on gene transcripts and proteins, as well as their biological significance. The phylogenetic tree construction tool bridges the gap between user-uploaded gene sequences of uncertain function and their homologs within the database’s 35 fish species. The comparative genome browser can provide phylogenetic relationships between fish species, as well as phylogenetic relationships and gene duplication events within specified gene families. In addition to the bulk transcriptome analysis, FishCODE’s cross-species transcriptome analysis tool enables the user to select samples across species for analysis. This may be particularly useful in understanding the biological processes that are shared across species, such as those involved in the immune response to viral attack (refer to “Cross-species transcriptome analysis” in Supplementary Document). Time series transcriptome analysis enables the utilization of different time points of data to elucidate the dynamics of an organism’s intrinsic regulatory processes, thereby identifying pivotal time frames and pivotal genes. Additionally, FishCODE’s epigenome analysis facilitates the execution of differential methylation region analysis, which offers a visual representation of the differential regions and associated genetic information.

Considering that most fish databases include transcriptome related analysis functions, for comparison, we also illustrate the capabilities and advantages of FishCODE analysis using its transcriptome analysis feature as an example. We focus on the time-series transcriptome analysis of the zebrafish retinal photodamage repair dataset (PRJNA748446) to elucidate the process. Users navigate to this dataset via species and experimental condition filters, the time points are divided according to the experimental meta-information of the sample provided by FishCODE (Fig. S9), with a cursory examination of the sample distribution and the application of preset analysis parameters, users can initiate the temporal transcriptome analysis with a click on the “Run” button.

According to the visualization results (Fig. 1G), users can obtain the time points corresponding to different levels e.g. level1 corresponds to control group, level4 corresponds to 5dpl (days post lesion) group. Notably, our analysis suggests that the level4 (5dpl) marks a significant transition in the gene expression network’s density, as revealed by the transcription factor (TF) gene coexpression network map across levels (Fig. 1H). The enrichment analysis using TF genes contained in level4 (5dpl) are shown in Figure (Fig. 1I). KEGG enriched many pathways related to post-injury repair, such as dre03083 (Polycomb repressive complex), dre04310 (Wnt signaling pathway), dre04350 (TGF-beta signaling pathway), and injury stress pathways dre04916 (Melanogenesis), dre04218 (Cellular senescence), dre05168 (Herpes simplex virus 1 infection), etc., suggesting that zebrafish photoinjury 5dpl at the time of photodamage or at the primary stage of post-damage repair. These findings are underpinned by experimental evidence (Fig. S10) (Kramer et al., 2021), which documents the nascent recovery of photoreceptor morphology in zebrafish retinae at this time point, underscoring its significance in the repair process. This insight enables the prioritization of genes from significantly enriched pathways for in-depth investigation into the molecular underpinnings of light-induced retinal damage repair. By analyzing this example dataset, researchers investigating damage repair mechanisms can complete the exploratory analysis of published genomics data in related fields in a one-stop manner without programming experience, and obtain information on the critical time points of light damage repair in zebrafish retina, systematic biological pathway changes, and the list of candidate genes at critical time points.

Paralleling the transcriptome analysis, FishCODE’s genomic and epigenomic analyses use the wealth of available omics data. Figure S5B shows differentially methylated regions in the liver between male and female carp and their associated genes. Figure S5C presents the phylogenetic relationship between the unnamed genes of coelacanth (*Latimeria chalumnae*) and highly similar genes from FishCODE. It suggests that the closer genetic distance between coelacanth and lungfish and the potential biological function of the unnamed gene may be related to the tp53 gene, corroborated by evolutionary insights from TimeTree (Kumar et al., 2022) and gene prediction algorithms from NCBI. FishCODE also offers a helpful information module for comprehensive usage details, including API interfaces (Fig. S11) and a dataset browsing module for user convenience and data access (Fig. S12).

In conclusion, FishCODE serves as a comprehensive multi-omics data information platform for fish, providing ample data resources and convenience for utilizing omics data in scientific research across diverse fields. FishCODE is an extensive and highly expandable system and will be maintained over time by specialized staff to ensure proper functioning and updates. Currently, the types of omics data encompassed by FishCODE are limited, and the methodologies for the integrated analysis of multi-omics data are still in their early stages. To enhance the relevance and utility of FishCODE as a research tool for scientists working with omics data, we plan to broaden its scope to include additional omics data types, specifically proteomics and metabolomics. Additionally, we aim to implement more effective integration methods for high-dimensional omics data, including advanced machine learning techniques.

## Data availability

The FishCODE database is publicly accessible through the website at http://bioinfo.ihb.ac.cn/fishcode. All data can be download at “DATASETS” Page of the website.

## Code availability

All the code used for omics data processing, online analysis, and visualization in FishCODE are freely available on https://github.com/hliihbcas/FishCODE. These codes can also be downloaded from the “OTHER TOOLS -> Omics preprocessing” menu on the FishCODE website.

## Acknowledgements

This work was supported by the National Key R&D Program of China [2021YFD1200804 and 2018YFD0901201] and the Strategic Priority Research Program of the Chinese Academy of Sciences (Precision Seed Design and Breeding) [XDA24010206]. We thank Ms. Zhixian Qiao from the Analysis and Testing Center at Institute of Hydrobiology for her technical support. The computational work in this study was supported by the Wuhan Branch, Supercomputing Center, Chinese Academy of Sciences, China.

